# Two Ume6 target gene classes in the biofilm regulatory network of *Candida albicans*

**DOI:** 10.64898/2026.09.24.753443

**Authors:** Eunsoo Do, C. Joel McManus, Aaron P. Mitchell

## Abstract

Biofilm formation by the fungal pathogen *Candida albicans* is a major source of infection. The biofilm regulator Ume6 forms complexes with partner transcription factors Efg1, Ndt80, and Upc2 to drive expression of biofilm-related genes. Here we present chromatin immunoprecipitation with sequencing (ChIP-seq) data for Ume6-chromatin association in both wild-type and *efg1*Δ/Δ *ndt80*Δ/Δ *upc2*Δ/Δ backgrounds. Ume6 associates with two promoter region classes. For one class, Ume6 association is significantly decreased in the *efg1*Δ/Δ *ndt80*Δ/Δ *upc2*Δ/Δ background. This class includes 577 genes, many with well-established roles in biofilm formation or the related process of filamentation. For the second class, Ume6 association is unperturbed in the triple mutant background. This class includes 534 genes, some with roles in iron homeostasis (e.g., *SFU1*) and ergosterol synthesis (e.g., *ERG11*), which may contribute to the impact of Ume6 on *ndt80*Δ/Δ mutant azole drug sensitivity. It also includes *FLO9*, a putative adhesin gene that is required for the emergent biofilm state produced by *efg1*Δ/Δ *ndt80*Δ/Δ double mutants. The data show that Ume6 binding to ∼half of its targets requires known partners, and Ume6 binding to ∼half of its targets does not. Ume6 may bind to the latter set of promoter regions independently of any partner, or perhaps with additional partners that have yet to be discovered.

**IMPORTANCE:** Biofilm formation has a central role in infection by the fungal pathogen *Candida albicans.*Here we focus on a key regulator, Ume6, that can drive biofilm formation. It is known to form protein complexes with three partner transcription factors, and here we investigate Ume6-promoter binding in the presence and absence of those partners. Ume6 binds to two promoter classes: one depends on known partners, and many associated genes have roles in biofilm formation. The second class is independent of known partners, and some associated genes are functionally connected to iron metabolism and ergosterol synthesis. The second class may mediate effects of Ume6 on azole drug sensitivity. Our results suggest that Ume6 may bind to some promoter regions independently of any partner, or perhaps with additional partners that have yet to be discovered.

## INTRODUCTION

Most microbes can colonize surfaces and thus grow as a biofilm (1). Microbial biofilms occur in nature, and have been exploited for a range of industrial applications (1, 2). However, biofilms pose a severe risk for human health when they form on tissues or implanted medical devices (1–3). In this situation, biofilm growth enables an infecting population to multiply, with risk compounded by the elevated drug tolerance that is typical of biofilms.

Our focus is *Candida albicans* (3), a fungal pathobiont that causes a spectrum of biofilm-related infections (3–6). Biofilm growth of *C. albicans* coincides with enormous changes in gene expression compared to its free-living planktonic state (7–10). Differentially expressed genes include several associated with filamentous growth and hypoxic adaption, two processes required for biofilm formation under many conditions. Although filamentous growth and hypoxic adaptation are distinct processes, they are coregulated in part through the transcription factors Efg1 and Ndt80 (11–14). In addition, numerous hypoxic adaptation genes depend upon the transcription factor Upc2 (15, 16). We recently described a distinct mechanism that couples filamentation and hypoxic adaptation: the transcription factor Ume6 interacts with Efg1, Ndt80, and Upc2 (12). At a genome-wide level, we found that loss of Efg1 reduces Ume6-chromatin association at several Ume6-activated genes (12). Here we present the impact of loss of all three partners – Efg1, Ndt80, and Upc2 – on Ume6-chromatin association.

## MATERIALS AND METHODS

### Strains and media

Strains used in this study were maintained in 15% glycerol frozen stocks at -80°C. Prior to use, cells were routinely grown on YPD agar plates (2% dextrose, 2% Bacto peptone, 1% yeast extract, 2% Bacto agar) for overnight at 30°C and then cultured in liquid YPD medium overnight at 30°C with agitation. Transformants were selected on YPD plus 400 μg/mL nourseothricin (clonNAT; Gold Biotechnology) or complete synthetic medium (2% dextrose, 1.7% Difco yeast nitrogen base with ammonium sulfate and auxotrophic supplements). All strains used in this study are listed in Table S1 in the supplemental material.

### Strain constructions

To manipulate *C. albicans* genome, the transient CRISPR-Cas9 system was employed as previously described in detail (17). Generally, the Cas9 cassette was amplified from the plasmid pV1093, and each of sgRNA cassette was generated by using split-joint PCR with “sgRNA/F” and “SNR52/R” as previously described in detail (17, 18). Primers and plasmids used for transformation are listed in Table S1.

To construct a *efg1*Δ/Δ *ndt80*Δ/Δ *upc2*Δ/Δ triple mutant strain in the *efg1*Δ/Δ *ndt80*Δ/Δ double mutant background, the *UPC2* deletion cassette was amplified from the plasmid pSFS2A-CaKan with primers ‘Upc2_dc_Kan/F’ and ‘Upc2_dc_Kan/R’ (19).

Transformants were screened on YPD containing 600 μg/ml G418 and 1.75 mg/ml molybdate, and candidates were genotyped by PCR using primers ‘Upc2 check up/F’ and ‘Upc2 check int/R’ for absence of *UPC2* ORF and using primers ‘Upc2 check up/F’ and ‘Kan int/R’ for the presence of the *CaKAN* marker at the *UPC2* locus.

To construct *UME6* overexpression strains (*P_RBT5_-UME6-HA*) in both the *efg1*Δ/Δ *ndt80*Δ/Δ double and the *efg1*Δ/Δ *ndt80*Δ/Δ *upc2*Δ/Δ triple mutant strains, the *NAT1-P_RBT5_* repair template and sgRNA cassette were prepared as previously described in detail (20). Then, the *UME6-3xHA-ADH1* terminator cassette was amplified from the plasmid “pSN52-HA-ADH1term” with primers “Ume6_F-HA” and “ADH1term -> HygB up/R”. The *CaHygB* cassette was amplified from the plasmid “pSFS2A-CaHygB” with primers “Kan_HygB/F” and “HygB -> Ume6 down/R”. Transformants were screened on YPD containing 600 μg/ml hygromycine B and 1.75 mg/ml quinine, and candidates were genotyped by PCR using primers ‘Ume6-tag confirm F’ and ‘Ume6-tag confirm R’ for absence of Ume6 terminator sequence and using primers ‘Kan_Hygb/F’ and ‘Ume6_3UTR_R’ for the presence of the *CaHYGB* marker at the *UME6* locus.

### ChIP-seq and Bioinformatic analysis

ChIP samples and ChIP-seq libraries were prepared according to previously described methods (21). Bioinformatic analysis used in this study was performed according to previously described methods (21). Differential binding analysis (DiffBind, Galaxy Version 2.10.0+galaxy0) (22), de novo motif discovery (HOMER, v 4.11) (23), ComputeMatrix (Galaxy Version 3.5.1.0.0), and PlotHeatmap (Galaxy Version 3.5.4+galaxy0) were conducted according to previously described methods (24). Gene Ontology (GO) term enrichment analysis was determined with the GO Termfinder tool at the *Candida* Genome Database (25).

### Data analysis software

ChIP-seq data were visualized using the Integrative Genomics Viewer v2.17.4 (26). Biofilm and filamentation images were processed using Image J (Fiji) (27). Statistical analyses and graph generations were carried out using GraphPad Prism version 9 (GraphPad Software, Inc., La Jolla).

### Data availability

Processed ChIP-seq data is available in Supplementary Table S2; raw data is available through NCBI SRA with accession numbers PRJNA1522041. Strains are available upon request.

## RESULTS AND DISCUSSION

For this study, we generated chromatin immunoprecipitation with sequencing (ChIP-Seq) data for a wild-type *C. albicans* strain and a derived *efg1*Δ/Δ *ndt80*Δ/Δ *upc2*Δ/Δ triple mutant. Each strain expressed a tagged *UME6-HA* allele or a control untagged *UME6* allele, which were fused to the *RBT5* promoter to circumvent Efg1- and Ndt80-dependence of *UME6* expression [as in previous studies (12)]. Strains were grown at 37°C for 4 hours in RPMI with 10% FBS, a low-iron medium that induces the *RBT5* promoter (12, 20).

We detected 1111 Ume6-bound genes in the wild-type *RBT5-UME6-HA* strain (Table S2). Binding of Ume6 to 577 genes was significantly reduced in the *efg1*Δ/Δ *ndt80*Δ/Δ *upc2*Δ/Δ triple mutant compared to the wild-type strain; we call these Efg1/Ndt80/Upc2-dependent genes (Fig 1A). In contrast, binding of Ume6 to 534 genes was not affected in the triple mutant compared to the wild-type strain; we call these Efg1/Ndt80/Upc2-independent genes (Fig 1B). These results indicate that only ∼half of Ume6-chromatin binding sites depend upon the known Ume6 partners Efg1, Ndt80, and Upc2.

**Figure 1.**
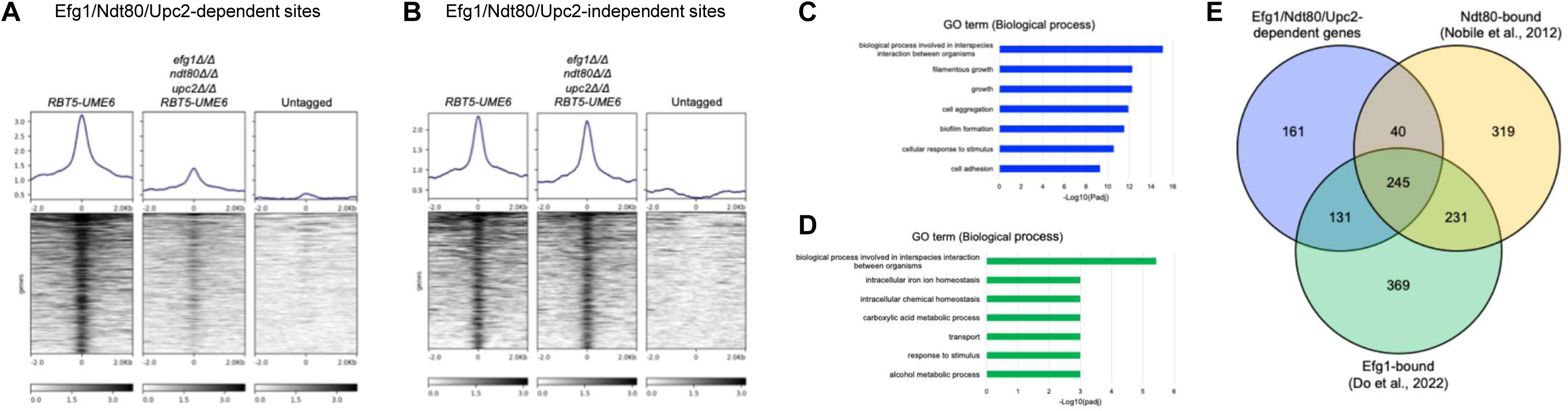
Two classes of Ume6-bound sites. (A and B) Heatmaps that display Ume6 binding intensities across strains. All strains were grown in RPMI + 10% FBS at 37°C for 4 hours, then used for ChIP-seq analysis. Ume6 DNA-binding at 577 Efg1/Ndt80/Upc2-dependent sites (A) or 534 Efg1/Ndt80/Upc2-independent sites (B) were compared between indicated strains. Each heatmap represents three biological replicates (n = 3). (C and D) Bar graph showing enriched GO terms from genes associated with the 577 Efg1/Ndt80/Upc2-dependent sites (C) or 534 Efg1/Ndt80/Upc2-independent sites (D). genes that show significantly abolished Ume6 binding sites in the *efg1*Δ/Δ *ndt80*Δ/Δ *upc2*Δ/Δ *RBT5-UME6* strain compared to the otherwise wild-type *RBT5-UME6* strain. (E) Venn diagram depicting the intersection of the Efg1/Ndt80/Upc2-dependent genes, Efg1-bound genes (21), and Ndt80-bound genes (10). Strains used in this study are listed in Supplementary file Table S1. Media composition is in Supplementary file Text S1. Standard methods were employed for strain construction (17–20) as detailed in Supplementary file Text S1. Primers and plasmids (17, 19, 21, 36–38) used for transformation are listed in Supplementary file Table S1. ChIP-seq experimental procedures and bioinformatic analysis were as previously described (21, 22, 25, 26) and detailed in Supplementary file Text S1.

The 577 Efg1/Ndt80/Upc2-dependent genes have several noteworthy features. The gene set is enriched for Gene Ontology (GO) terms related to interspecies interaction, filamentation and biofilm formation (Fig 1C). We found that 285 of the 577 genes are direct Ndt80 targets (10) (Fig 1E). In addition, 376 of the 577 genes are direct Efg1 targets (12) (Fig 1E), and 245 of the 577 genes are targets of both Efg1 and Ndt80.

Many genes among these Efg1/Ndt80 direct targets have been well studied for roles in biofilm formation or filamentation. Examples include adhesin genes *ALS1* and *ALS3*; toxin gene *ECE1*; hyphal polarity gene *HGC1*; and filamentation regulatory genes *WOR3, ROB1, NRG1, EFG1,* and *BCR1*. Only 26 of the 577 genes are direct Upc2 targets (28) (Fig S1). Although the number of Upc2-bound genes is small, several have roles in biofilm formation or hypoxic adaptation, including *ACE2, ERG10, ERG2, ERG251,* and *ERG6*. Therefore, functions of many Efg1/Ndt80/Upc2-dependent genes align with the functions of Ume6, Efg1, Ndt80, and Upc2.

The contributions of Efg1, Ndt80, and Upc2 to Ume6-chromatin binding is reflected in motif enrichment within the 577 genes. Our de novo analysis revealed enrichment for an Ndt80-like motif (ACACAAA, p-value= 1e-40), a Upc2-like motif (TCGTCT, p-value= 1e-30), and an Efg1-like motif (TGCAT, p-value= 1e-15) among the Efg1/Ndt80/Upc2-dependent sites (Fig S2). These same motifs were evident in strongly bound Ume6 sites in our previous study (12). Enrichment for these motifs is consistent with the finding that Ume6 binding to these targets is Efg1/Ndt80/Upc2-dependent.

For the 577 Efg1/Ndt80/Upc2-dependent genes, we found previously that an *efg1*Δ/Δ mutation reduced Ume6 binding at many of these promoters (12). *WOR3, BCR1,* and *ALS3* are examples in which Ume6 binding was decreased in an *efg1*Δ/Δ background, but is more severely decreased in the triple mutant background (Fig 2A). We suggest that Efg1 is the main partner that promotes Ume6 binding at these promoters. *CLB2* is an example in which Ume6 binding was unaffected in an *efg1*Δ/Δ background, but is severely decreased in the triple mutant background (Fig 2A). We suggest that either Efg1 or Ndt80 can promote Ume6 binding at the *CLB2* promoter, or perhaps Ndt80 is the major contributor to Ume6 binding. *ERG251* is bound by Upc2 but not Efg1 or Ndt80. For this gene, Ume6 binding was unaffected in an *efg1*Δ/Δ background, but is decreased in the triple mutant background (Fig 2A). We suggest that Upc2 is the main partner that promotes Ume6 binding at this promoter.

**Figure 2.**
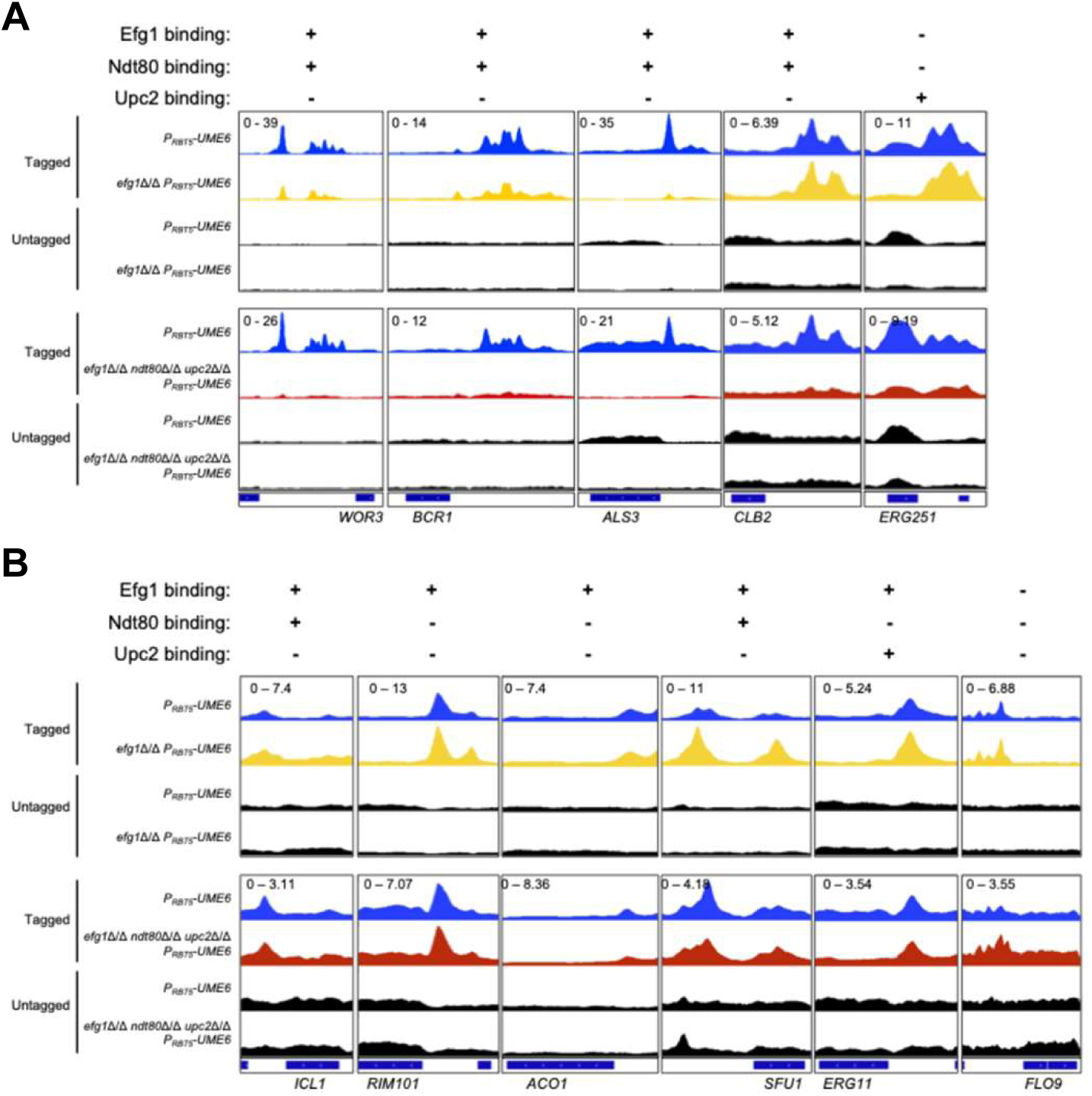
Illustrative Ume6-bound genomic regions. (A and B) Genome browser tracks that show Ume6 binding in upstream regions of indicated genes. The Y axis indicates read counts. Data from Do et al. (12) are shown in the upper strip of panels; data from the current study are shown in the lower strip. Tracks of untagged strains were used as controls. All ChIP-seq tracks represent three biological replicates (n = 3). Genomic regions include those upstream of (A) Efg1/Ndt80/Upc2-dependent genes *WOR3, BCR1, ALS3, CLB2,* and *ERG251;* (B) Efg1/Ndt80/Upc2-independent genes *ICL1, RIM101, ACO1, SFU1, ERG11,* and *FLO9.* Assignments of binding by Efg1, Ndt80, and Upc2 come from published studies (10, 12, 28).

The 534 Efg1/Ndt80/Upc2-independent genes also have several noteworthy features. GO analysis of the 534 genes showed enrichment for interspecies interaction (Fig 1D), with representative genes that include *ICL1* and *RIM101* (Fig 2B). GO terms related to iron homeostasis were also enriched (Fig 1D). In fact, iron- and Ume6-regulated genes such as *ACO1* and *SFU1* (Fig. 2B), which may contribute to the impact of Ume6 on azole drug sensitivity of *ndt80*Δ/Δ mutants (29), were in this gene set. The ergosterol biosynthetic genes in this set, such as *ERG11* (Fig 2B), may also contribute to the azole sensitivity phenotype (29). Finally, we note that the putative adhesin gene *FLO9* is included in this gene set (Fig. 2B). *FLO9* is activated by Ume6 in an *efg1*Δ/Δ *ndt80*Δ/Δ double mutant and is required for the double mutant’s emergent biofilm phenotype (24). We would predict that Ume6 binding to the *FLO9* promoter region would be independent of Efg1 and Ndt80, a prediction supported by this dataset.

We determined de novo binding motifs for Ume6 binding sites of the 534 Efg1/Ndt80/Upc2-independent genes. These sites lacked Ndt80- or Efg1-like motifs, but we identified an enriched Upc2-like motif: TCGTCT (P-value: 1e-76). This motif can be recognized more broadly by zinc-cluster transcription factors (30). Its enrichment raises the possibility that Ume6 alone, or in complexes with other zinc-cluster regulators, may bind to that motif (Fig S3).

The features of Ume6-bound genomic regions align well with our understanding of Ume6 function. Ume6 associates with regulatory regions of numerous genes tied to filamentation and biofilm formation, the major biological functions deduced for Ume6 by Kadosh and colleagues (31–35). Ume6 also associates with regulatory regions of ergosterol and iron homeostasis genes, which may explain the impact of Ume6 on fluconazole sensitivity (29). For many of these regulatory regions, Ume6 binding is impaired by loss of Efg1, Ndt80, and Upc2, the three transcription factors with which Ume6 is known to interact (12). Our results also open up new questions about Ume6 function. Ume6 binds to the regulatory regions of yeast-associated genes such as *NRG1* and *YWP1*; does Ume6 function in those contexts as a repressor? Also, about half of Ume6-bound regions are unaffected by loss of the three known Ume6 partners; does Ume6 have additional partners that direct its binding to these regions? Finally, we detect Ume6 bound to iron homeostasis regulatory genes *SFU1* and *RIM101*, and to carbon metabolic genes *ICL1* and *ACO1*; does Ume6 have a global impact on carbon or heme regulation? We hope that future researchers discover the answers to these questions.

## ACKNOWLEDGMENTS

We are grateful to Drs. Katharina Goerlich, Anupam Sharma, Yinhe Mao, Liping Xiong, Min-Ju Kim, Max Cravener, Amelia White, and Fred Lanni for their continued interest and suggestions. We thank Max Kuhr for outstanding laboratory management.

---

**Fig S1.**
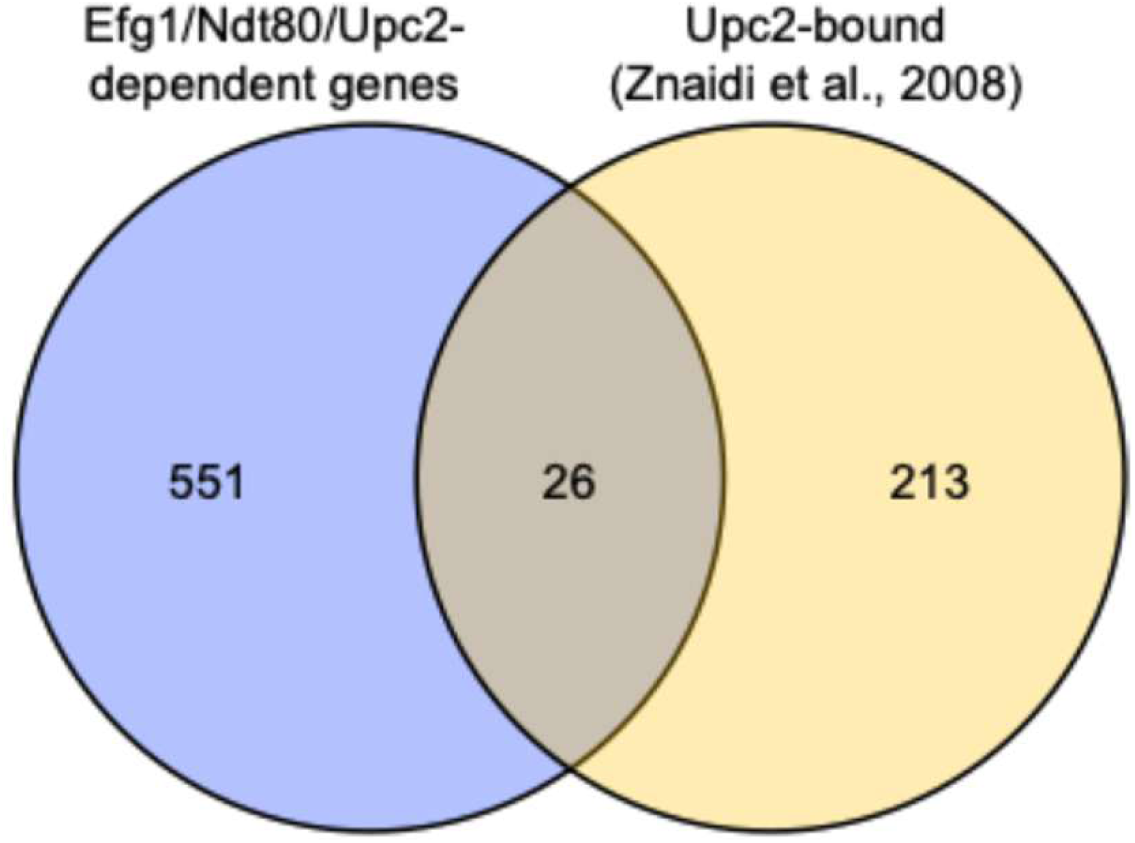
Upc2-bound genes in the Efg1/Ndt80/Upc2-dependent genes. Venn diagram depicts intersection of the Efg1/Ndt80/Upc2-dependent genes and the Upc2-bound genes (Znaidi et al., 2008).

**Fig S2.**
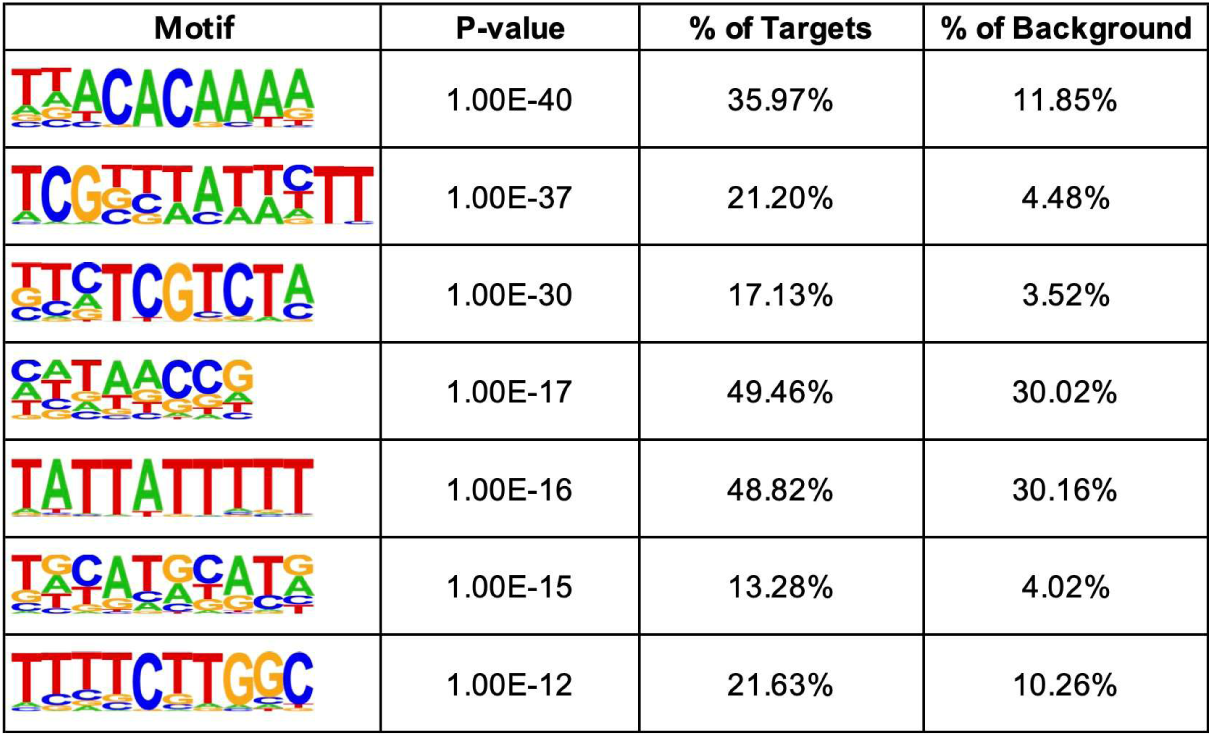
Enriched binding motifs in the Efg1/Ndt80/Upc2-dependent genes. *De novo* binding motif analysis for the 577 gene promoters was performed with HOMER.

**Fig S3.**
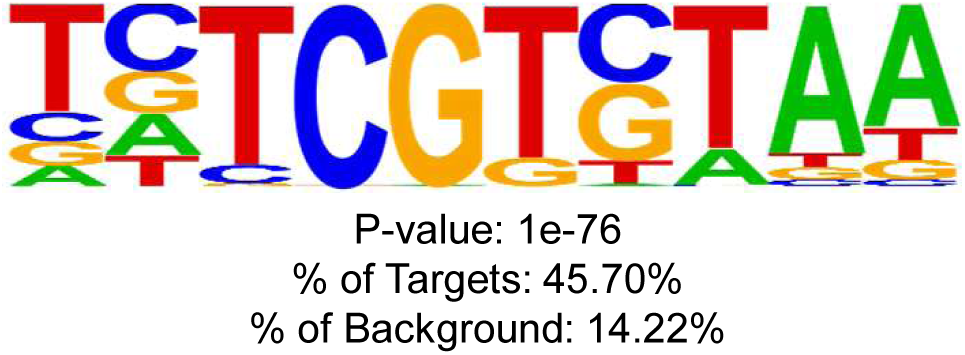
Enriched binding motif in the Efg1/Ndt80/Upc2-independent genes. *De novo* binding motif analysis for the 534 gene promoters was performed with HOMER.

